# Negative Impact of Daily Screen Use on Inhibitory Control Network in Preadolescence: A Two-Year Follow-Up Study

**DOI:** 10.1101/2022.02.22.481547

**Authors:** Ya-Yun Chen, Hyungwook Yim, Tae-Ho Lee

## Abstract

The COVID-19 pandemic has made an unprecedented shift in children’s daily lives. Children are increasingly spending time with screens to learn and connect with others. As the online environment rapidly substitutes in-person experience, understanding children’s neuropsychological trajectories associated with screen experiences is important. Previous findings suggest that excessive screen use can lead children to prefer more immediate rewards over delayed outcomes. We hypothesized that increased screen time delays a child’s development of inhibitory control. By analyzing neuropsychological data from 8,324 children (9-11ys) from the ABCD Study, we found that children who had more screen time showed a higher reward orientation and a weaker inhibitory control system (i.e., fronto-striatal circuitry) in the brain. Importantly, we found that the interaction between screen exposure and reward sensitivity negatively influenced the development of the inhibitory control system in the brain over a two year period. These results indicate possible negative long-term impacts of increased daily screen time on children’s neuropsychological development. The results further demonstrated that screen time influences dorsal striatum connectivity, which suggests that the effect of daily screen use is a habitual seeking behavior. The study provides neural and behavioral evidence on the negative impact of daily screen use on developing children.

## Introduction

Childhood is a critical period for the development of inhibitory control (IC) (Williams et al., 1999), where the ability to resist impulsive behaviors (Carlson et al., 2002; Duckworth & Kern, 2011) and focus on long-term goals changes significantly along with their related neural circuits (Padmanabhan et al., 2011). Studies have demonstrated the negative impact of excessive screen exposure on the development of children’s inhibitory control (Carson et al., 2016; Domingues-Montanari, 2017; Twenge & Campbell, 2018) by highlighting the nature of the screen platform, which often offers immediate benefits with negligible costs (Frey et al., 2007). For example, most screen platforms allow users to pause and skip any session that they are less interested in and to choose contents that please them more. This increased information accessibility allows children to pursue immediate rewards and feedback (Tricomi & Fiez, 2012), and, in turn, this increased reward-seeking tendency can further weaken the development of IC (Burton et al., 2021).

Although social organizations and governments offer guidelines about limiting screen time for school-age children to mitigate its negative impact (Communications et al., 2016; Okely et al., 2019), the average use of screen time entertainment for children is continuously increasing (Tsiros et al., 2017; Twenge & Campbell, 2018), and has been amplified since the COVID-19 pandemic. For example, screen exposure time has been escalated by at least 50% as virtual experiences substitute in-person interactions during the period of the Stay-at-Home order (SuperAwesome, 2020).

Recent evidence suggests that the underlying neural mechanisms linked to the failure of IC may be associated with the imbalance between the executive networks and the regions involved in reward-related processing (e.g., amygdala, striatum, ventral tegmental area) (Casey, 2015; Lee & Telzer, 2016; McClure et al., 2004; Metcalfe & Mischel, 1999). Although non-clinical daily screen behaviors are understudied, research in screen dependency disorders (SDD) with various age groups (from children aged 5-11, adolescents aged 9-17, to college-aged young adults) shows that the reward-related nucleus and the central executive networks which may be affected by daily screen behaviors are the striatum and the frontoparietal network (FPN) (Balleine et al., 2007; Brand et al., 2014).

The striatum is composed of three nuclei (i.e., caudate, putamen, and ventral striatum), which integrate input from the brainstem and a variety of subcortical and cortical regions, and is involved in motor planning, decision-making, motivation, and reward perception and responses (Taylor et al., 2013; Yager et al., 2015). Both animal and human studies have revealed that the striatum serves a key role in habit formation and addictive processes (Corbit et al., 2012; Everitt et al., 2008; Everitt & Robbins, 2013; Schwendt et al., 2009; Volkow et al., 2006). For example, in rodents, long access to self-administered methamphetamine gradually decreased dopamine transporter protein levels in the striatum (Schwendt et al., 2009). Similarly, a human imaging study using positron emission tomography (PET) revealed that the metabolism in the striatum was decreased when the individual with cocaine addiction saw the cocaine cue compared with a neutral cue, where the magnitude of the reduction positively correlated with their craving (Volkow et al., 2006). A task-based functional MRI (fMRI) showed that young adult patients with internet addiction exhibited less activation in the striatum compared to controls when they continuously won in a competitive task (Dong et al., 2013). Regarding volumetric comparison, increased striatal volume was found in young adult smokers compared to nonsmokers (Li et al., 2015) and in young adults with SDD (Cai et al., 2016). Developmentally, adolescents who excessively play computer games have been reported to have higher gray matter volume and higher activation in the striatum compared to those who play computer games infrequently (Kühn et al., 2011). These findings suggest that the striatum is associated with reward processing and screen dependency.

The FPN, primarily comprised of the dorsolateral prefrontal cortex (DLPFC) and the posterior parietal cortex (Uddin et al., 2019), and its DLPFC node were investigated extensively in the context of inhibitory control and dependency behaviors since it is involved in executive and regulatory processes in the brain (Brand et al., 2014; Hare et al., 2009; Hayashi et al., 2013; Lopez et al., 2019; Vincent et al., 2008). For example, Lopez and colleagues (2019) used an elegant two-session study to reveal that the dieters that had to recruit greater DLPFC activation in the first IC exertion session showed less FPN activity in the subsequence food-cue task and tended to consume more ice cream when the diets were broken (Lopez et al., 2019). Studies with screen dependency disorders (SDD) also reported that young adults with SDD have dysfunctions in the DLPFC and the FPN during resting-state and task-based scans (Dong et al., 2015; Liu et al., 2014). During the inhibitory task, the SDD group exhibited a lower inhibition efficiency with lower activation and connectivity in the FPN regulatory regions compared to the control group, which suggests that the inhibitory function was impaired in the disorder group. Consistently, by using the resting-state functional connectivity approach, studies also reported that the SDD group showed more within-network connectivity reduction in FPN compared to the control group (Dong et al., 2015). They further reported that the strength of the connectivity was negatively correlated with the Stroop effect. Taken together, these studies suggest that the function of the FPN is crucial for one’s control ability during reward-related processing and is also related to screen use.

Although the critical role of a certain brain region, such as the FPN, DLPFC, or striatum, in inhibitory control processing has been investigated extensively as a single region or network, recent studies highlight the importance of neural connectivity in inhibitory processing at the large-scale brain systems level. In particular, recent neural circuitry research from both animal and human studies suggests that it is more comprehensive to consider the between-level neural connectivity as a system (Figure 1A). The FPN and the striatum can also be viewed from a brain-systems level as they are structurally and functionally interconnected, being part of the fronto-striatal circuit, subserving inhibitory control through the DLPFC node (Alexander et al., 1986; Haber, 2016; Leh et al., 2007; Rubia et al., 2006; Sigman, 2017; Zhang & Iwaki, 2020). For example, functional MRI (fMRI) studies using diverse task-based indices demonstrated a progressive maturation for IC functions within the fronto-striatal circuitry, where the level of activation in the circuitry increased along with inhibition efficiency across development (Rubia et al., 2006). The findings suggest that the neural maturation of IC can be examined by measuring functional coupling between the FPN and the striatum, and that the regions should be considered together when investigating IC.

**Figure 1.**
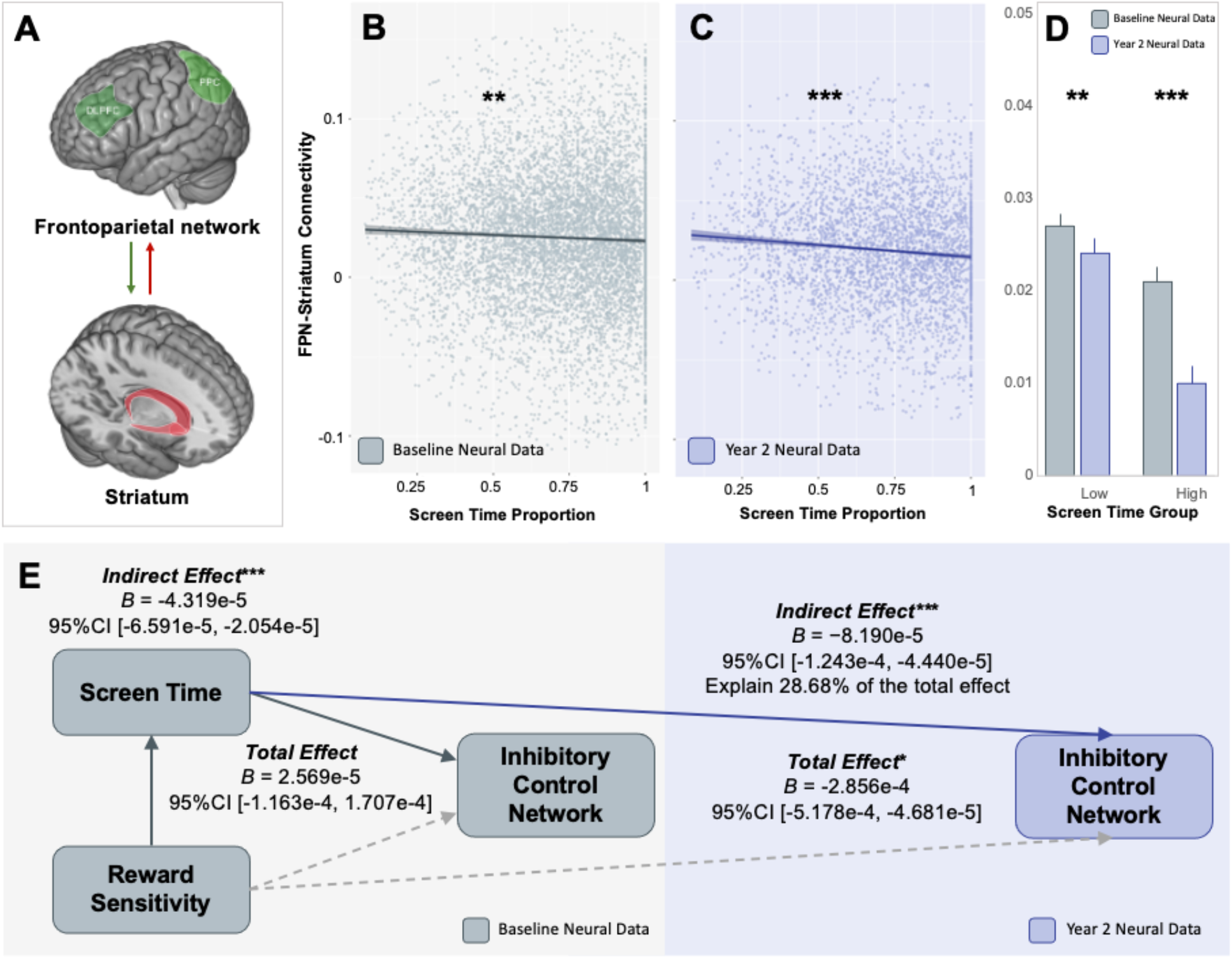
Association between screen exposure, brain, and reward sensitivity. **A:** Illustrations of the ROIs examined. **B:** Correlation between the baseline year Screen-Activity Proportion (SAP) and the FPN-Striatum Connectivity from the baseline year neural data. **C:** Correlation between the baseline year SAP and the FPN-Striatum Connectivity from the year 2 neural data. **D:** The group comparisons between high SAP and low SAP in FPN-Striatum Connectivity for both years. **E:** Association between reward sensitivity (BAS score), screen time (SAP), and the inhibitory control network (FPN-Striatum Connectivity) from the baseline data (in grey) and the year 2 data (in blue). * p < .05 or BF_10_ > 3;** p < .01 or BF_10_ > 10; *** p < .001 or BF_10_ > 30.

Given previous evidence indicating that the maturation of the brain can be indexed by the degree of functional coupling between the FPN and the striatum (Balleine et al., 2007; Brand et al., 2014; Casey, 2015; Liston et al., 2006; Rubia et al., 2006), the main goal of the present study is to delineate how daily screen exposure time influences the development of the intrinsic neurocircuitry in children that underlies IC (i.e., fronto-stratal circuitry, FPN-striatum). We used the baseline and post-baseline year 2 follow-up waves of the Adolescent Brain Cognitive Develpment study (Casey et al., 2018) to examine the intrinsic functional connectivity estimated from resting-state fMRI, focusing on the functional connectivity between the frontoparietal network and the striatum. We hypothesized that (1) there would be an alteration of the fronto-striatal connectivity in children with longer daily screen exposure time, and (2) the daily screen exposure time in the baseline year (year of enrollment) can predict the fronto-striatal connectivity strength in year 2 (i.e., post-baseline year 2 follow-up) as well as the strength change between waves. In addition to the effect of daily screen exposure time in the inhibitory control network, we further predicted that (3) as children become more exposed to screens, their reward sensitivity will be augmented, which will delay the development of the inhibitory neural circuit (Johnson et al., 2003; Kim et al., 2016; Kim-Spoon et al., 2016; Miller et al., 2004).

## Materials and Methods

### Participants and data preprocessing

The present study was performed by using data retrieved from the Adolescent Brain Cognitive Development (ABCD) Data Repository (https://abcdstudy.org). The ABCD Data Repository tracks 11,878 individuals aged 9 to 11 years from 21 data collection sites around the United States (Casey et al., 2018). The dataset was downloaded on 11/2020 from the ABCD 2.0 release from the NIH Data Archive (NDA, https://nda.nih.gov/). The present study included the ABCD Youth Screen Time Survey (STQ, *abcd_stq01*), the Sports and Activities Survey (*abcd_spacss01*), the ABCD Youth Behavioral Inhibition/Behavioral Approach System Scales (*abcd_bisbas01*) and the MRI-related measures include the processed resting-state fMRI data (ABCD rsfMRI Network to Subcortical ROI Correlations, *mrirscor02*). After removing the missing value, the final sample size was 8,324 for the baseline year (interview date from 09/2016 to 10/2018) and 3,891 for the year 2 follow-up (interview date from 07/2018 to 01/2020). The age range of children in this sample in the baseline year was 108-131 months old (*M* = 119.28, *SD* = 7.47) and 49.65% of them were female. Children were primarily identified as White (76.99%), 18.69% as African American, and 4.32% as Others. Caregiver ages ranged from 23 to 80 years (*M* = 40.21, *SD* = 6.75). See Table S1 for more demographic information.

### Demographic and behavioral data

#### Demographics Survey

In a parent-report questionnaire, caregivers reported the race and gender of themselves and their child, family structure, socioeconomic status (SES, total household income), education, and religion.

#### Screen Time Assessment

Screen time was assessed using a child self-report measure consisting of 14 items (Barch et al., 2018; Sharif et al., 2010). The main 12 items measured different types of screen utilization, such as watching TV shows or movies, watching videos, and playing video games, on a typical weekday and weekend day. The scale is as follows: 0 = None; .25 = < 30 minutes; 0.5 = 30 minutes; 1 = 1 hour; 2 = 2 hours; 3 = 3 hours; and 4 = 4+ hours. Additionally, the survey included two items related to the experiences in playing mature-rated video games and watching R-rated movies. Daily average screen time was calculated by averaging the sum of weekday screen time and the sum of weekend time.

#### Non-Screen Activities Time Assessment

Activities time refers to all sports and activities other than screen time, which was measured by the parent-reported Sports and Activities Involvement Questionnaire (Huppertz et al., 2016). The items in the questionnaire included 31 different types of sports, music, art, and hobbies. Parents had to report how many years, how many months per year, how many days per week, and how many minutes per session their child participated in a certain activity. The activities time data was processed in two steps: missing value remedy and final score calculation.

The data set had several missing values for various types of activity which were dispersed amongst participants. When discarding individuals with missing values, 2,075 samples (25%) had to be removed. Thus the treatment rules that were applied were: 1) for missing values of days per week items, take out all data points that have the same value on months per year and then calculate the median of days per week of those data points for replacing the missing values. For example, if soccer: days per week is missing and the months per year is 2, then all other months per year that are 2 under soccer will be taken out and then the median of days per week of them will be calculated. The median value will be used to replace the missing values in this example. 2) for missing values of minutes per session, take out all data points that have the same value of months per year and then calculate the median of minutes per session of those data points for replacing the missing point.

After treating the missing values, since the scale of the survey was different from the scale of the screen time survey, daily average activity time was calculated by the following steps: 1) convert the minutes per session of a single type of activity to hour per session; 2) multiply the hour per session and days per week of a single type of activity and divided by seven; 3) add up all types of activities.

#### Screen-Activity Proportion Score

In the examinations, the non-screen activities time was used to control the screen exposure time by the following calculation:

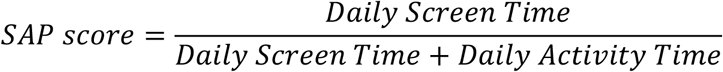

The Screen-Activity Proportion (SAP) score reflects how much individuals are exposed to the screen on a daily basis compared to non-screen activities. A SAP score above 0.5 means that an individual spends more time daily with screens than on non-screen activities.

#### Behavioral Inhibition/Behavioral Activation System (BIS/BAS) Scales

Carver and White (Carver & White, 1994) suggested two motivation systems, the behavioral inhibition system (BIS) and the behavioral activation system (BAS). The BIS corresponds to the motivation to avoid aversive outcomes. The BAS corresponds to the motivation to approach reward outcomes, which can be allocated to three facets: BAS Drive, BAS Fun-seeking, and BAS Reward Responsiveness. To examine the relationship between individual behavioral traits in reward sensitivity, the degree of screen time exposure, and neural development, we included BAS scores, provided by the ABCD dataset. Although the BAS is divided into three subscales, it was found to load onto one single dimension significantly when a factor analysis was conducted (Miller et al., 2004). Thus, in the current analysis, we used the sum of the three BAS subscales (i.e., BAS Total), as an indication of overall reward sensitivity (Kelley et al., 2019).

### Neuroimaging data

#### Imaging protocol

The ABCD imaging protocol was integrated from three 3T scanner platforms (i.e., Siemens Prisma [66.87%], General Electric/GE 750 [20.48%], and Philips [18.06%]), which used standard adult-size multi-channel coils that are capable of multiband echo-planar imaging (EPI) acquisitions with repetition time (TR) = 0.8 s, voxel size = 2.4 mm isotropic, 60 slices, field of view (FoV) = 216×216 mm, echo time (TE) = 30 ms, flip angle = 52 degrees. The twenty minutes resting-state data was acquired with eyes open and passive viewing of a crosshair. Briefly, the resting-state data was processed with motion correction, B0 distortion correction, resampled to isotropic, rigidly coregistered to the structural image, and spatially normalized. More information about the protocols, such as scanning parameters and motion correction processing, are available in Casey et al. (2018) and Hagler et al. (2019).

#### Resting-State Intrinsic connectivity between the Inhibitory Control Network to Subcortical ROI connectivity

The ABCD dataset provides processed resting-state intrinsic network connectivity with various subcortical regions based on the subcortical segmentation with the FreeSurfer’s automated brain segmentation (aseg) atlas (Bruce, 2012) to the cortical networks’ that were extracted by the Gordon parcellation approach (Gordon et al., 2016). For the present study, the values representing the strength of the frontoparietal network (FPN) to striatum connectivity (Figure 1A) were generated by averaging the scores of the FPN-bilateral caudate, the FPN-bilateral putamen, and the FPN-bilateral accumbens connectivities. More details about the MRI processing pipeline and ROI extraction are available in Hagler et al. (Hagler Jr et al., 2019).

### Data analysis

Because of the heterogeneous nature of the neural data of the ABCD study, the effect sizes represented by *r* of the current study are small. Several ABCD neural correlate studies were reported with a small effect size as the current study (Cheng et al., 2020; Karcher et al., 2019; Paulus et al., 2019; Rosenberg et al., 2020). To ensure the robustness of the results, the current study applied a Bayesian approach and a Bootstrap Hypothesis Testing (Dick et al., 2021) to ensure the robustness of the results.

Data analyses were performed by combining a 50,000 permutation resampling method and the robust method (Pernet et al., 2013), unless otherwise noted. By performing the resampling with replacement (bootstrapping and permutation) techniques, ordinary least squares (OLS) models can be used without meeting assumptions of normality and homoscedasticity (Efron & Tibshirani, 1994; Manly, 2006). The statistical results from both Frequentist (i.e., *p*-value) and Bayesian approaches were reported. Bayesian approaches are able to evaluate the relative plausibility of the alternative hypothesis (H_1_) and the null hypothesis (H_0_) at the same time, which can avoid the inflated false positive rate due to the large sample size (i.e., more than 10,000 in the ABCD data; c.f., (Sullivan & Feinn, 2012). We used Bayes factors (BF_10_) as support for the alternative hypothesis (H_1_) over the null hypothesis (H_0_). A BF_10_ larger than 3 is interpreted as moderate favor for H_1_ and a value larger than 10 is interpreted as a strong favor for H_1_ (Matzke, 2014). For the Bayesian test, because the previous ABCD neural correlate studies reported significant results with a small effect size (*r*’s range from 0.037 to 0.07) (Cheng et al., 2020; Karcher et al., 2019; Paulus et al., 2019; Rosenberg et al., 2020), in the present study, a conservative stretched beta prior width of 0.3 was set, reflecting the belief of a medium effect size.

## Results

### Association between screen exposure and fronto-striatal connectivity

The correlation analysis showed a significant negative association between the screen-activity proportion and the fronto-striatal connectivity of the baseline year (*r* = -0.040, 95% *CI*^*50,000*^ bootstrap = [-0.0605, -0.0186], *BF*_*10*_ = 13.10). As the strength of the fronto-striatal connectivity reflects the maturation quality of the inhibitory control system (Rubia et al., 2006), the results suggest that a longer screen exposure time was significantly related to the underdevelopment of the inhibitory control system in the brain (Figure 1B).

An additional group-level comparison with children who had a Screen-Activity Proportion (SAP) greater than the 90th percentile (SAP = 1, *N* = 785) and children who had an SAP less than the 10th percentile (SAP < 0.39, *N* = 788, *M*_*SAP*_ = 0.29, *SD* = 0.07), confirmed that children with longer screen time showed a significantly lower fronto-striatal connectivity, compared to children with shorter screen time (Figure 1D in grey; *M*_*high*_ = 0.021, *M*_*low*_ = 0.027, independent *t*-test: *t*_*(1571)*_ = 3.043, *p* = 0.002, Cohen’s *d* = 0.153, *BF*_*10*_ = 5.487).

### Association between screen exposure and the development of fronto-striatal connectivity

A correlation analysis using the year 2 neural data demonstrated a stronger significant negative association between the screen-activity proportion and the fronto-striatal connectivity than with the baseline neural data (*r* = -0.0933, 95% *CI*^*50,000*^ bootstrap = [-0.1231, -0.0635], *BF*_*10*_ = 428995.74; Figure 1C). In addition, daily screen exposure time for the baseline year predicted the connectivity strength change between baseline and the second year (year 2 minus baseline) (*r* = -0.0340, 95% *CI*^*50,000*^ bootstrap = [-0.0645, -0.0035], *BF*_*10*_ = 0.349).

As conducted in the baseline data, there was a statistically significant difference in fronto-striatal connectivity between the high SAP group (*N* = 329) and the low SAP group (*N* = 382, *M*_*SAP*_ = 0.28, *SD* = 0.08) in the year 2 data. The group effect confirmed that the longer screen activity is associated with the development of the inhibitory control system in the brain (Figure 1D in blue; *M*_*high*_ = 0.010, *M*_*low*_ = 0.024, *t*_*(709)*_ = 5.675, *p* < 0.001, Cohen’s *d* = 0.427, *BF*_*10*_ = 439537.31).

Along with the result of the baseline neural correlates, the results of the developmental data suggest that a longer screen exposure time may negatively affect the development of the inhibitory control system in the brain.

### Association between reward sensitivity, screen exposure, and brain

The pairwise correlations between reward sensitivity trait, screen exposure, and inhibitory control (*Table S4*) imply that screen exposure time may mediate the relationship between trait reward sensitivity and inhibitory control. Therefore, we further hypothesized that screen exposure time mediates the relationship between the BAS score and the strength of the fronto-striatal connectivity.

A bootstrapping mediation analysis was used (Biesanz et al., 2010; Preacher & Hayes, 2008) to test the hypothesis that screen-activity proportion mediates the relationship between the BAS score and the strength of the fronto-striatal connectivity. The results showed that the screen-activity proportion significantly mediated the effect between the BAS score and the year 2 fronto-striatal connectivity (indirect effect: *B* = −8.190e-5, *SE* = 1.981e-5, *p* < 0.001, 95% *CI*^*5,000*^ bootstrap = [-1.243e-4, -4.440e-5]; total effect: *B* = -2.856e-4, *SE* = 1.150e-4, *p* < 0.05, 95% *CI*^*5,000*^ bootstrap = [-5.178e-4, - 4.681e-5]), explaining 28.68% of the total effect. This mediation effect was also shown when using the baseline fronto-striatal connectivity (indirect effect: *B* = -4.319e-5, *SE* = 1.229e-5, *p* < 0.001, 95% *CI*^*5,000*^ bootstrap = [-6.591e-5 -2.054e-5]; total effect: *B* = 2.569e-5, *SE* = 7.196e-5, *p* = 0.721, 95% *CI*^*5,000*^ bootstrap = [-1.163e-4 1.707e-4]). The results suggest that the screen exposure effect augments the aversive reward sensitivity effect on the inhibitory control network development. The results with nuisance regressors (sex, age, household income, and parental education level) are shown in Table S2. We also evaluated the model with BAS as a mediator, which yielded a non-significant result (See *Supplemental Materials Section D* for more details).

### Association between screen exposure and the subdivisions of the striatum

Previous studies suggest that the ventral striatum (i.e., nucleus accumbens, Nacc) is more related to simple voluntary behaviors whereas the dorsal striatum (i.e., caudate and putamen) plays a major role in habitual seeking and addictive behaviors (Everitt & Robbins, 2013; Zhou et al., 2018). Thus, we additionally hypothesized that the longer daily screen exposure time leads children to the more habitual seeking behavior associated with increased dorsal striatum activation. To test this hypothesis, we also examined the screen exposure effect for each striatal subregion and found that the longer screen exposure was more associated with the decreased dorsal striatum-FPN connectivity, especially in the year 2, *r*_*caudate*_ = -0.0597, 95% *CI*^*50,000*^ = [-0.0902, -0.0292], *BF*_*10,caudate*_ = 15.738; *r*_*putamen*_ = -0.0710, 95% *CI*^*50,000*^ = [-0.1010, -0.0414], *BF*_*10,putamen*_ = 245.716). In contrast, there was no such association with the ventral striatum (*r*_*Nacc*_ = - 0.0257, 95% *CI*^*50,000*^ = [-0.0560, 0.0047], *BF*_*10,Nacc*_ = 0.070. (See *Supplemental Materials Section E* for more details).

### Validating the association between intrinsic FPN-striatum connectivity and the function of inhibitory control

The current study adopted FPN-striatum to represent the intrinsic neurocircuitry that underlies IC. We utilized several behavioral indices of inhibitory function, including the behavioral performances in the Flanker Inhibitory Control and Attention Test (Eriksen & Eriksen, 1974), the Monetary Incentive Delay Task (Everitt et al., 2008), the Stop Signal Task (Logan, 1994), ADHD trait (Achenbach & Edelbrock, 1991), and the Cash Choice Task. For the task descriptions and details of results see Supplemental Materials Section G. The results suggest that the resting state FPN-Striatum connectivity and the effect of screen exposure were able to anchor to the behavioral indices of inhibitory function.

## Discussion

In our two-year follow-up study, we examined how daily screen exposure time can influence the neural inhibitory control network (ICN) in children. The results demonstrated that a longer screen exposure time was negatively associated with the strength of the fronto-striatal circuitry. The results further demonstrated that a longer screen exposure time predicts a protracted development of the ICN supported by a negative correlation with the strength of the fronto-striatal connectivity in year 2 and the strength change between the two waves. Finally, the results showed that a longer screen exposure mediated the effect between the reward sensitivity and the wave2 fronto-striatal connectivity, indicating that a longer screen exposure may augment the aversive reward sensitivity effect on ICN development. Importantly, longer screen exposure was associated with the dorsal striatum, which is involved in habitual seeking behavior and addictive behavior.

Our findings on the inverse fronto-striatal connectivity in both cross-sectional and longitudinal data extend previous work by demonstrating that the screen exposure effect is not linked only to a single region, but can influence the interaction between regions and systems (Balleine et al., 2007; Brand et al., 2014). Previous studies using fMRI and DTI suggest that increased positive functional connectivity between the striatum and the frontal executive network is associated with enhanced efficiency of IC across development (Liston et al., 2006; Rubia et al., 2006). Thus, negative fronto-striatal connectivity among children who have higher screen exposure may reflect decreasing sophistication of functional coupling between these networks as the brain matures. This novel finding is supported by studies on substance addiction. For example, both DTI and functional connectivity techniques revealed a negative functional coupling of the fronto-striatal connectivity during smoking cue-induced craving (Yuan et al., 2017) and in alcohol-dependent patients relative to healthy controls (Becker et al., 2017). Similarly, a higher daily screen exposure can weaken the development of the fronto-striatal network, which is supported by our findings which show that fronto-striatal connectivity changes in individuals with increased screen dependency.

Consistent with the previous findings (Becker et al., 2017; Kohno et al., 2014; Motzkin et al., 2014; Wang et al., 2013; Yuan et al., 2017), we found a detrimental effect of the negative fronto-striatal coupling in inhibitory process. However, it should be noted that there is contradictory evidence showing that longer exposure to addictive stimuli was associated with positive fronto-striatal connectivity (Hu et al., 2015; Koehler et al., 2013). For example, a recent resting-state fMRI study showed that pathological gambling patients have increased connectivity between ventral striatum and the superior/middle frontal gyrus (Koehler et al., 2013), and cocaine addiction patients showed an increased fronto-striatal connectivity (Hu et al., 2015). Furthermore, it has been demonstrated that the negative coupling between frontal and limbic networks is associated with better self-control ability (Lee & Telzer, 2016). However, given the heterogeneity of study populations and approaches, these inconsistent findings in fronto-striatal coupling are not unexpected. Some of the diverse reports may be due to distinct addictive behaviors or the status of those behaviors, such as period of ‘at risk’, current dependence, and the period of recovery (Pariyadath et al., 2016). Moreover, the utilization of different target regions within fronto-striatal circuits may explain these contradictory findings. For instance, one study showed a positive coupling by using the ventral striatum, whereas another showed a negative coupling by employing the caudate and dorsal striatum (Wang et al., 2013).

Importantly, the current study showed a difference in the connectivity between each subdivision of the striatum and the frontal executive network. High screen exposure resulted in a more negative dorsal striatum coupling to the frontal executive network relative to the ventral striatum, especially in the second-wave data. Evidence from both rodent and human studies has shown a functional shifting from ventral to dorsal striatum underlying the voluntary behaviors becoming compulsion, suggesting a dysfunction of the IC (Everitt et al., 2008; Everitt & Robbins, 2013; Zhou et al., 2018). In the rodent model, as cocaine seeking became habitual for the rats, dopamine release was increased in the dorsal, but not ventral, striatum (Everitt et al., 2008). Similarly, a study with cannabis-dependent males showed a lower dorsal striatum connectivity in the frontal regions compared to the controls (Zhou et al., 2018). Therefore, it is possible that the consequence of long-term extensive screen exposure may be similar to the effect of the addictive behaviors (Dong et al., 2021). Individuals who have excessive screen exposure induce frontal network connectivity shifting from the ventral to the dorsal striatum, leading them to be more vulnerable to impulses and addictions. However, due to an insufficient amount of substance dependency measurements in the currently released ABCD dataset, it is hardly possible to do a meaningful statistic. An explicit test of the inference is expected with the more released data or in other future studies.

We also exhibited the generalizability of the negative effect of screen exposure by controlling the sex and age of the children as well as their parental SES and education level. Previous studies showed that there are demographic differences in screen exposure in children and adolescents, for example boys reported higher overall screen use than girls (Nagata et al., 2022), and youth from lower SES and minority backgrounds engaged in more screen activity (Reid Chassiakos et al., 2016). Additionally, there is a demonstrated correlation between parental SES and both children’s executive function (Lawson et al., 2018) and the development of fronto-striatal circuitry during adolescence (Li et al., 2022). By accounting for these factors, the current study demonstrated that the delaying effect of screen exposure on the development of the ICN is a general phenomenon existing in children during preadolescence.

The current two-year follow-up study demonstrates that excessive daily screen exposure can shift functional connectivity patterns, suggesting that long-term screen exposure may affect the development of the ICN and can excessively augment the reward-seeking tendency. We used several behavioral indices of inhibitory function to validate that the intrinsic connectivity of ICN and the effect of screen exposure were associated with inhibitory control outcomes. However, we are limited by the currently released processed data in the ABCD study, as we have had difficulty in gaining the processed inhibitory task-based cortical network to subcortical regions data, which would allow us to observe how this altered intrinsic ICN works during inhibitory tasks. In future studies it will be necessary to examine how daily screen exposure influences connectivity in the fronto-striatal network while the inhibitory processing is activating.

In summary, we showed that with longer daily screen exposure, neural coupling between the frontoparietal network and striatum decreases in children. Importantly, we also found that this excessive daily screen exposure may augment a child’s reward-seeking tendency and the consequence of the intrinsic neural connectivity is similar to impulsive addictive behaviors. Given that the virtual movement is irreversible and expanding in the modern and future life, we appeal for increasing research attention to daily screen exposure.

## ACKNOWLEDGMENTS

This work was supported by the Virginia Tech ISCE research to T.-H. Lee and the National Research Foundation of Korea funded by the Ministry of Science and ICT (No.2018R1A5A7059549) to H. Yim.

Data used in the preparation of this article was obtained from the Adolescent Brain Cognitive Development (ABCD) Study (https://abcdstudy.org), held in the NIMH Data Archive (NDA). This is a multisite, longitudinal study designed to recruit more than 10,000 children aged 9-10 and follow them over 10 years into early adulthood. The ABCD Study® is supported by the National Institutes of Health and additional federal partners under award numbers U01DA041048, U01DA050989, U01DA051016, U01DA041022, U01DA051018, U01DA051037, U01DA050987, U01DA041174, U01DA041106, U01DA041117, U01DA041028, U01DA041134, U01DA050988, U01DA051039, U01DA041156, U01DA041025, U01DA041120, U01DA051038, U01DA041148, U01DA041093, U01DA041089, U24DA041123, U24DA041147. A full list of supporters is available at https://abcdstudy.org/federal-partners.html. A listing of participating sites and a complete listing of the study investigators can be found at https://abcdstudy.org/consortium_members/. ABCD consortium investigators designed and implemented the study and/or provided data but did not necessarily participate in the analysis or writing of this report. This manuscript reflects the views of the authors and may not reflect the opinions or views of the NIH or ABCD consortium investigators. The ABCD data repository grows and changes over time. The ABCD data used in this report came from ABCD Annual Release 2.0. DOIs can be found at https://dx.doi.org/10.15154/1503209.

## DATA AVAILABILITY

Source Data: https://osf.io/z9dcp/?view_only=bb229c96747a4213ad36ada37651ec28

## Reference

Achenbach, T. M., & Edelbrock, C. (1991). Child behavior checklist. Burlington (Vt), 7, 371–392.

Alexander, G. E., DeLong, M. R., & Strick, P. L. (1986). Parallel organization of functionally segregated circuits linking basal ganglia and cortex. Annual review of neuroscience, 9(1), 357–381.

Balleine, B. W., Delgado, M. R., & Hikosaka, O. (2007). The role of the dorsal striatum in reward and decision-making. Journal of Neuroscience, 27(31), 8161–8165.

Barch, D. M., Albaugh, M. D., Avenevoli, S., Chang, L., Clark, D. B., Glantz, M. D., … Yurgelun-Todd, D. (2018). Demographic, physical and mental health assessments in the adolescent brain and cognitive development study: Rationale and description. Developmental cognitive neuroscience, 32, 55–66.

Becker, A., Kirsch, M., Gerchen, M. F., Kiefer, F., & Kirsch, P. (2017). Striatal activation and frontostriatal connectivity during non-drug reward anticipation in alcohol dependence. Addiction biology, 22(3), 833–843.

Biesanz, J. C., Falk, C. F., & Savalei, V. (2010). Assessing mediational models: Testing and interval estimation for indirect effects. Multivariate Behavioral Research, 45(4), 661–701.

Brand, M., Young, K. S., & Laier, C. (2014). Prefrontal control and Internet addiction: a theoretical model and review of neuropsychological and neuroimaging findings. Frontiers in human neuroscience, 8, 375.

Bruce, F. (2012). Freesurfer. Neuroimage, 62(2), 774–781.

Burton, S., Knibb, G., & Jones, A. (2021). A meta-analytic investigation of the role of reward on inhibitory control. Quarterly Journal of Experimental Psychology, 17470218211008895.

Cai, C., Yuan, K., Yin, J., Feng, D., Bi, Y., Li, Y., … Tian, J. (2016). Striatum morphometry is associated with cognitive control deficits and symptom severity in internet gaming disorder. Brain imaging and behavior, 10(1), 12–20.

Carlson, S. M., Moses, L. J., & Breton, C. (2002). How specific is the relation between executive function and theory of mind? Contributions of inhibitory control and working memory. Infant and Child Development: An International Journal of Research and Practice, 11(2), 73–92.

Carson, V., Hunter, S., Kuzik, N., Gray, C. E., Poitras, V. J., Chaput, J.-P., … Connor Gorber, S. (2016). Systematic review of sedentary behaviour and health indicators in school-aged children and youth: an update. Applied physiology, nutrition, and metabolism, 41(6), S240–S265.

Carver, C. S., & White, T. L. (1994). Behavioral inhibition, behavioral activation, and affective responses to impending reward and punishment: the BIS/BAS scales. Journal of personality and social psychology, 67(2), 319.

Casey, B. (2015). Beyond simple models of self-control to circuit-based accounts of adolescent behavior. Annual review of psychology, 66, 295–319.

Casey, B., Cannonier, T., Conley, M. I., Cohen, A. O., Barch, D. M., Heitzeg, M. M., … Garavan, H. (2018). The adolescent brain cognitive development (ABCD) study: imaging acquisition across 21 sites. Developmental cognitive neuroscience, 32, 43–54.

Cheng, W., Rolls, E., Gong, W., Du, J., Zhang, J., Zhang, X.-Y., … Feng, J. (2020). Sleep duration, brain structure, and psychiatric and cognitive problems in children. Molecular psychiatry, 1–12.

Communications, C. o., Media, & MBE. (2016). Media use in school-aged children and adolescents. Pediatrics, 138(5), e20162592.

Corbit, L. H., Nie, H., & Janak, P. H. (2012). Habitual alcohol seeking: time course and the contribution of subregions of the dorsal striatum. Biological psychiatry, 72(5), 389–395.

Dick, A. S., Lopez, D. A., Watts, A. L., Heeringa, S., Reuter, C., Bartsch, H., … Marshall, A. (2021). Meaningful associations in the adolescent brain cognitive development study. Neuroimage, 118262.

Domingues-Montanari, S. (2017). Clinical and psychological effects of excessive screen time on children. Journal of paediatrics and child health, 53(4), 333–338.

Dong, G., Hu, Y., Lin, X., & Lu, Q. (2013). What makes Internet addicts continue playing online even when faced by severe negative consequences? Possible explanations from an fMRI study. Biological psychology, 94(2), 282–289.

Dong, G., Lin, X., & Potenza, M. N. (2015). Decreased functional connectivity in an executive control network is related to impaired executive function in Internet gaming disorder. Progress in Neuro-Psychopharmacology and Biological Psychiatry, 57, 76–85.

Dong, G.-H., Dong, H., Wang, M., Zhang, J., Zhou, W., Du, X., & Potenza, M. N. (2021). Dorsal and ventral striatal functional connectivity shifts play a potential role in internet gaming disorder. Communications biology, 4(1), 1–9.

Duckworth, A. L., & Kern, M. L. (2011). A meta-analysis of the convergent validity of self-control measures. Journal of research in personality, 45(3), 259–268.

Efron, B., & Tibshirani, R. J. (1994). An introduction to the bootstrap. CRC press.

Eriksen, B. A., & Eriksen, C. W. (1974). Effects of noise letters upon the identification of a target letter in a nonsearch task. Perception & psychophysics, 16(1), 143–149.

Everitt, B. J., Belin, D., Economidou, D., Pelloux, Y., Dalley, J. W., & Robbins, T. W. (2008). Neural mechanisms underlying the vulnerability to develop compulsive drug-seeking habits and addiction. Philosophical Transactions of the Royal Society B: Biological Sciences, 363(1507), 3125–3135.

Everitt, B. J., & Robbins, T. W. (2013). From the ventral to the dorsal striatum: devolving views of their roles in drug addiction. Neuroscience & Biobehavioral Reviews, 37(9), 1946–1954.

Frey, B. S., Benesch, C., & Stutzer, A. (2007). Does watching TV make us happy? Journal of Economic psychology, 28(3), 283–313.

Gordon, E. M., Laumann, T. O., Adeyemo, B., Huckins, J. F., Kelley, W. M., & Petersen, S. E. (2016). Generation and evaluation of a cortical area parcellation from resting-state correlations. Cerebral Cortex, 26(1), 288–303.

Haber, S. N. (2016). Corticostriatal circuitry. Dialogues in clinical neuroscience, 18(1), 7.

Hagler Jr, D. J., Hatton, S., Cornejo, M. D., Makowski, C., Fair, D. A., Dick, A. S., … Harms, M. P. (2019). Image processing and analysis methods for the Adolescent Brain Cognitive Development Study. Neuroimage, 202, 116091.

Hare, T. A., Camerer, C. F., & Rangel, A. (2009). Self-control in decision-making involves modulation of the vmPFC valuation system. Science, 324(5927), 646–648.

Hayashi, T., Ko, J. H., Strafella, A. P., & Dagher, A. (2013). Dorsolateral prefrontal and orbitofrontal cortex interactions during self-control of cigarette craving. Proceedings of the National Academy of Sciences, 110(11), 4422–4427.

Hu, Y., Salmeron, B. J., Gu, H., Stein, E. A., & Yang, Y. (2015). Impaired functional connectivity within and between frontostriatal circuits and its association with compulsive drug use and trait impulsivity in cocaine addiction. JAMA psychiatry, 72(6), 584–592.

Huppertz, C., Bartels, M., de Zeeuw, E. L., van Beijsterveldt, C. E., Hudziak, J. J., Willemsen, G., … de Geus, E. J. (2016). Individual differences in exercise behavior: stability and change in genetic and environmental determinants from age 7 to 18. Behavior Genetics, 46(5), 665–679.

Johnson, S. L., Turner, R. J., & Iwata, N. (2003). BIS/BAS levels and psychiatric disorder: An epidemiological study. Journal of psychopathology and behavioral assessment, 25(1), 25–36.

Karcher, N. R., O’Brien, K. J., Kandala, S., & Barch, D. M. (2019). Resting-state functional connectivity and psychotic-like experiences in childhood: results from the adolescent brain cognitive development study. Biological psychiatry, 86(1), 7–15.

Kelley, N. J., Kramer, A. M., Young, K. S., Echiverri-Cohen, A. M., Chat, I. K.-Y., Bookheimer, S. Y., … Zinbarg, R. E. (2019). Evidence for a general factor of behavioral activation system sensitivity. Journal of research in personality, 79, 30–39.

Kim, Y., Jeong, J.-E., Cho, H., Jung, D.-J., Kwak, M., Rho, M. J., … Choi, I. Y. (2016). Personality factors predicting smartphone addiction predisposition: Behavioral inhibition and activation systems, impulsivity, and self-control. PloS one, 11(8), e0159788.

Kim-Spoon, J., Deater-Deckard, K., Holmes, C., Lee, J., Chiu, P., & King-Casas, B. (2016). Behavioral and neural inhibitory control moderates the effects of reward sensitivity on adolescent substance use. Neuropsychologia, 91, 318–326.

Koehler, S., Ovadia-Caro, S., van der Meer, E., Villringer, A., Heinz, A., Romanczuk-Seiferth, N., & Margulies, D. S. (2013). Increased functional connectivity between prefrontal cortex and reward system in pathological gambling. PloS one, 8(12), e84565.

Kohno, M., Morales, A. M., Ghahremani, D. G., Hellemann, G., & London, E. D. (2014). Risky decision making, prefrontal cortex, and mesocorticolimbic functional connectivity in methamphetamine dependence. JAMA psychiatry, 71(7), 812–820.

Kühn, S., Romanowski, A., Schilling, C., Lorenz, R., Mörsen, C., Seiferth, N., … Büchel, C. (2011). The neural basis of video gaming. Translational psychiatry, 1(11), e53–e53.

Lawson, G. M., Hook, C. J., & Farah, M. J. (2018). A meta-analysis of the relationship between socioeconomic status and executive function performance among children. Developmental science, 21(2), e12529.

Lee, T.-H., & Telzer, E. H. (2016). Negative functional coupling between the right fronto-parietal and limbic resting state networks predicts increased self-control and later substance use onset in adolescence. Developmental cognitive neuroscience, 20, 35–42.

Leh, S. E., Ptito, A., Chakravarty, M. M., & Strafella, A. P. (2007). Fronto-striatal connections in the human brain: a probabilistic diffusion tractography study. Neuroscience letters, 419(2), 113–118.

Li, M., Lindenmuth, M., Tarnai, K., Lee, J., King-Casas, B., Kim-Spoon, J., & Deater-Deckard, K. (2022). Development of cognitive control during adolescence: The integrative effects of family socioeconomic status and parenting behaviors. Developmental cognitive neuroscience, 57, 101139.

Li, Y., Yuan, K., Cai, C., Feng, D., Yin, J., Bi, Y., … von Deneen, K. M. (2015). Reduced frontal cortical thickness and increased caudate volume within fronto-striatal circuits in young adult smokers. Drug and alcohol dependence, 151, 211–219.

Liston, C., Watts, R., Tottenham, N., Davidson, M. C., Niogi, S., Ulug, A. M., & Casey, B. (2006). Frontostriatal microstructure modulates efficient recruitment of cognitive control. Cerebral Cortex, 16(4), 553–560.

Liu, G.-C., Yen, J.-Y., Chen, C.-Y., Yen, C.-F., Chen, C.-S., Lin, W.-C., & Ko, C.-H. (2014). Brain activation for response inhibition under gaming cue distraction in internet gaming disorder. The Kaohsiung Journal of Medical Sciences, 30(1), 43–51.

Logan, G. D. (1994). On the ability to inhibit thought and action: A users’ guide to the stop signal paradigm.

Lopez, R. B., Courtney, A. L., & Wagner, D. D. (2019). Recruitment of cognitive control regions during effortful self-control is associated with altered brain activity in control and reward systems in dieters during subsequent exposure to food commercials. PeerJ, 7, e6550.

Manly, B. F. (2006). Randomization, bootstrap and Monte Carlo methods in biology (Vol. 70). CRC press.

Matzke, D. (2014). Bayesian explorations in mathematical psychology. Universiteit van Amsterdam [Host].

McClure, S. M., Laibson, D. I., Loewenstein, G., & Cohen, J. D. (2004). Separate neural systems value immediate and delayed monetary rewards. Science, 306(5695), 503–507.

Metcalfe, J., & Mischel, W. (1999). A hot/cool-system analysis of delay of gratification: dynamics of willpower. Psychological review, 106(1), 3.

Miller, E., Joseph, S., & Tudway, J. (2004). Assessing the component structure of four self-report measures of impulsivity. Personality and Individual differences, 37(2), 349–358.

Motzkin, J. C., Baskin-Sommers, A., Newman, J. P., Kiehl, K. A., & Koenigs, M. (2014). Neural correlates of substance abuse: reduced functional connectivity between areas underlying reward and cognitive control. Human brain mapping, 35(9), 4282–4292.

Nagata, J. M., Ganson, K. T., Iyer, P., Chu, J., Baker, F. C., Gabriel, K. P., … Bibbins-Domingo, K. (2022). Sociodemographic Correlates of Contemporary Screen Time Use among 9-and 10-Year-Old Children. The Journal of Pediatrics, 240, 213-220.e212.

Okely, A., Ghersi, D., Loughran, S., Cliff, D., Shilton, T., Jones, R., … Eckermann, S. (2019). Australian 24-hour movement guidelines for children and young people (5-17 years): An integration of physical activity, sedentary behaviour and sleep–research report. Australian Government Department of Health.

Padmanabhan, A., Geier, C. F., Ordaz, S. J., Teslovich, T., & Luna, B. (2011). Developmental changes in brain function underlying the influence of reward processing on inhibitory control. Developmental cognitive neuroscience, 1(4), 517–529.

Pariyadath, V., Gowin, J. L., & Stein, E. A. (2016). Resting state functional connectivity analysis for addiction medicine: from individual loci to complex networks. Progress in brain research, 224, 155–173.

Paulus, M. P., Squeglia, L. M., Bagot, K., Jacobus, J., Kuplicki, R., Breslin, F. J., … Bartsch, H. (2019). Screen media activity and brain structure in youth: evidence for diverse structural correlation networks from the ABCD study. Neuroimage, 185, 140–153.

Pernet, C. R., Wilcox, R., & Rousselet, G. A. (2013). Robust correlation analyses: false positive and power validation using a new open source matlab toolbox. Frontiers in psychology, 3, 606.

Preacher, K. J., & Hayes, A. F. (2008). Asymptotic and resampling strategies for assessing and comparing indirect effects in multiple mediator models. Behavior research methods, 40(3), 879–891.

Reid Chassiakos, Y. L., Radesky, J., Christakis, D., Moreno, M. A., Cross, C., Hill, D., … Boyd, R. (2016). Children and adolescents and digital media. Pediatrics, 138(5).

Rosenberg, M. D., Martinez, S. A., Rapuano, K. M., Conley, M. I., Cohen, A. O., Cornejo, M. D., … Wager, T. D. (2020). Behavioral and neural signatures of working memory in childhood. Journal of Neuroscience, 40(26), 5090–5104.

Rubia, K., Smith, A. B., Woolley, J., Nosarti, C., Heyman, I., Taylor, E., & Brammer, M. (2006). Progressive increase of frontostriatal brain activation from childhood to adulthood during event-related tasks of cognitive control. Human brain mapping, 27(12), 973–993.

Schwendt, M., Rocha, A., See, R. E., Pacchioni, A. M., McGinty, J. F., & Kalivas, P. W. (2009). Extended methamphetamine self-administration in rats results in a selective reduction of dopamine transporter levels in the prefrontal cortex and dorsal striatum not accompanied by marked monoaminergic depletion. Journal of Pharmacology and Experimental Therapeutics, 331(2), 555–562.

Sharif, I., Wills, T. A., & Sargent, J. D. (2010). Effect of visual media use on school performance: a prospective study. Journal of Adolescent Health, 46(1), 52–61.

Sigman, A. (2017). Screen Dependency Disorders.

Sullivan, G. M., & Feinn, R. (2012). Using effect size—or why the P value is not enough. Journal of graduate medical education, 4(3), 279–282.

SuperAwesome. (2020). Everyone is a kids and family brand now: Data, observations and recommendations for companies interacting with kids and families during lockdown. https://www.superawesome.com/everyone-is-a-kids-and-family-brand-now/

Taylor, S. B., Lewis, C. R., & Olive, M. F. (2013). The neurocircuitry of illicit psychostimulant addiction: acute and chronic effects in humans. Substance abuse and rehabilitation, 4, 29.

Tricomi, E., & Fiez, J. A. (2012). Information content and reward processing in the human striatum during performance of a declarative memory task. Cognitive, Affective, & Behavioral Neuroscience, 12(2), 361–372.

Tsiros, M. D., Samaras, M. G., Coates, A. M., & Olds, T. (2017). Use-of-time and health-related quality of life in 10-to 13-year-old children: not all screen time or physical activity minutes are the same. Quality of Life Research, 26(11), 3119–3129.

Twenge, J. M., & Campbell, W. K. (2018). Associations between screen time and lower psychological well-being among children and adolescents: Evidence from a population-based study. Preventive medicine reports, 12, 271–283.

Uddin, L. Q., Yeo, B., & Spreng, R. N. (2019). Towards a universal taxonomy of macro-scale functional human brain networks. Brain topography, 32(6), 926–942.

Vincent, J. L., Kahn, I., Snyder, A. Z., Raichle, M. E., & Buckner, R. L. (2008). Evidence for a frontoparietal control system revealed by intrinsic functional connectivity. Journal of neurophysiology, 100(6), 3328–3342.

Volkow, N. D., Wang, G.-J., Telang, F., Fowler, J. S., Logan, J., Childress, A.-R., … Wong, C. (2006). Cocaine cues and dopamine in dorsal striatum: mechanism of craving in cocaine addiction. Journal of Neuroscience, 26(24), 6583–6588.

Wang, Y., Zhu, J., Li, Q., Li, W., Wu, N., Zheng, Y., … Wang, W. (2013). Altered fronto-striatal and fronto-cerebellar circuits in heroin-dependent individuals: a resting-state FMRI study. PloS one, 8(3), e58098.

Williams, B. R., Ponesse, J. S., Schachar, R. J., Logan, G. D., & Tannock, R. (1999). Development of inhibitory control across the life span. Developmental psychology, 35(1), 205.

Yager, L. M., Garcia, A. F., Wunsch, A. M., & Ferguson, S. M. (2015). The ins and outs of the striatum: role in drug addiction. Neuroscience, 301, 529–541.

Yuan, K., Yu, D., Bi, Y., Wang, R., Li, M., Zhang, Y., … Lu, X. (2017). The left dorsolateral prefrontal cortex and caudate pathway: New evidence for cue-induced craving of smokers. Human brain mapping, 38(9), 4644–4656.

Zhang, F., & Iwaki, S. (2020). Correspondence Between Effective Connections in the Stop-Signal Task and Microstructural Correlations. Frontiers in human neuroscience, 14, 279.

Zhou, F., Zimmermann, K., Xin, F., Scheele, D., Dau, W., Banger, M., … Becker, B. (2018). Shifted balance of dorsal versus ventral striatal communication with frontal reward and regulatory regions in cannabis-dependent males. Human brain mapping, 39(12), 5062–5073.

