## Supplementary Information for "Negative Impact of Daily Screen Use on Inhibitory Control Network in Preadolescence: A Two-Year Follow-Up Study"

**Author Contributions:** Y.-Y. Chen developed the study concept. Y.-Y. Chen analyzed and interpreted data under the supervision of T.-H. Lee and H. Yim. Y.-Y. Chen drafted the manuscript, and T.-H. Lee and H. Yim provided critical revisions. All authors approved the final version of the manuscript for submission.

**Competing Interest Statement:** The authors declare no competing interests.

**Keywords:** screen time, inhibitory control, child development, fronto-striatal circuits, fronto-parietal network, striatum.

### A. Demographic variables

The demographic information, including sample size, interview date, female percentage, age (in month), caregiver's age (in year), race, caregiver's marital status, total combined family income, and parental education, for both waves were reported in **Table S1**.

***Table S1.*** *Demographic table*

| | Wave 1 | Wave 2 |
|---|---|---|
| Sample size (*N*) | 8,324 | 3,891 |
| Interview date | 09/2016-10/2018 | 07/2018-01/2020 |
| Female percentage | 49.64% | 46.94% |
| Age (in month) | 119.32 (107-133) | 143.49 (127-163) |
| Caregiver's age (in year) | 40.21 (23-80) | 42.35 (25-82) |
| **Race alone** | | *(from wave 1 data)* |
| White | 66.61% | 71.34% |
| African American | 13.69% | 10.80% |
| Native American[a] | 0.46% | 0.46% |
| Asian | 1.92% | 1.59% |
| Others | 3.99% | 3.50% |
| Multiracial population | 11.85% | 11.11% |
| **Caregiver's marital status** | | |
| Married | 70.16% | 71.21% |
| Living with partner | 5.22% | 5.76% |
| Others | 24.01% | 22.39% |
| **Total combined family income[b]** | | |
| < $25,000 | 11.51% | 8.30% |
| $25,000 - $49,999 | 12.79% | 11.03% |
| $50,000 - $99,999 | 26.33% | 26.25% |

| | | |
|---|---|---|
| $100,000 - $199,999 | 30.19% | 33.68% |
| >=$200,000 | 11.37% | 13.42% |
| Refuse to answer or don’t know | 7.83% | 7.30% |
| **Parental Education**[c] | | *(from wave 1 data)* |
| <High school diploma | 5.55% | 4.19% |
| High school Diploma or GED | 9.87% | 8.66% |
| Some college or associate degree | 28.23% | 28.89% |
| Bachelor's degree | 29.68% | 31.90% |
| Post graduate degree | 26.54% | 26.22% |
| Refuse to answer | 0.13% | 0.13% |

*All demographic variables were parent-reported. For example, for the race attribute, the question was: “What race do you consider the child to be?.”*

[a]*Native American included American Indian, Alaska Native, and Native Hawaiian.*

[b]What is your total combined highest income for the past 12 months?

[c]*What is the highest grade or level of school you have completed or the highest degree you have received?*

### B. Bayes Factor Robustness Check

With robustness check (**Figure S1**), we can see that the screen time effect on both the baseline and year 2 neural data results were robust ($BF_{10} > 3$). However, panel C shows the connectivity strength change between the baseline and the second year is in the anecdotal range ($BF_{10}$ = 0.33-3).

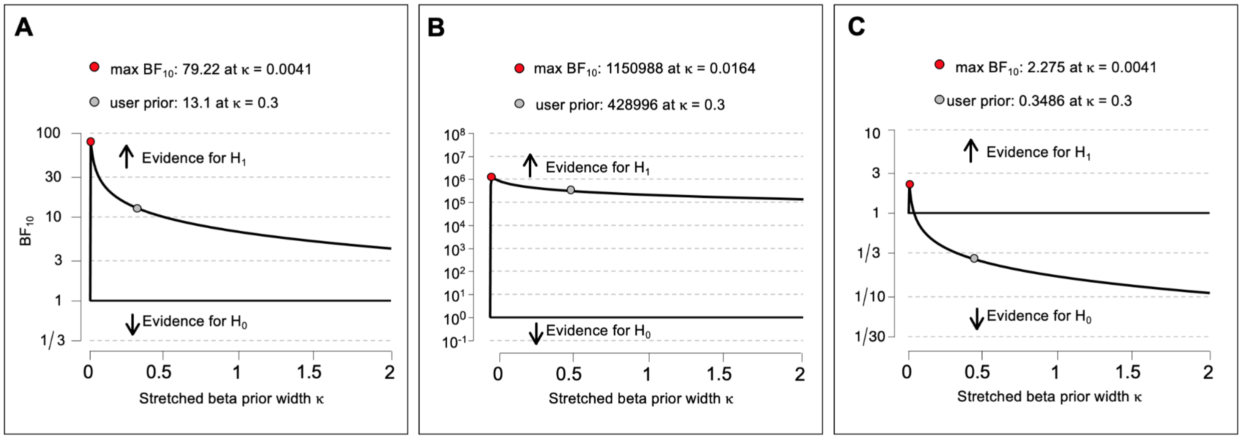


**Screen x FPN-Striatum (Wave1)** **Screen x FPN-Striatum (Wave2)** **Screen x FPN-Striatum (Wave2-Wave1)**

***Figure S1.*** *Robustness check graphs for three Bayesian examination.* ***A:*** *Robustness check for correlation between the baseline year Screen-Activity Proportion (SAP) and the FPN-Striatum Connectivity from the baseline neural data.* ***B:*** *Robustness check for correlation between the baseline year SAP and the FPN-Striatum Connectivity from the year 2 neural data.* ***C:*** *Robustness check for correlation between the baseline year SAP and the difference between the year 2 and baseline FPN-Striatum Connectivity.*

### C. The results of mediation analysis with nuisance regressors

The results (**Table S2**) suggest that the daily screen exposure effect augments the aversive reward sensitivity effect on the inhibitory control network development even with controlling the sex, age, and socioeconomic status. The main model indicates a mediation model with screen exposure as a mediator to mediate the effect of reward sensitivity on the brain. The panels following the main model show the results with controlling the effects from different demographic confounding variables.

***Table S2.*** *The results of Mediation analysis*

| | ***B*** | ***SE*** | ***p*** | ***95% CI$^{5,000}$ bootstrap*** |
|---|---|---|---|---|
| **Main model (Baseline year)** | | | | |
| Indirect effect | -4.319e-5 | 1.229e-5 | < 0.001 | [-6.591e-5 -2.054e-5] |
| Total effect | 2.569e-5 | 7.196e-5 | 0.721 | [-1.163e-4 1.707e-4] |
| *Controlling for sex and age* | | | | |
| Indirect effect | -4.436e-5 | 1.264e-5 | < 0.001 | [-6.932e-5 -2.120e-5] |
| Total effect | 1.909e-5 | 7.516e-5 | 0.800 | [-1.228e-4 1.723e-4] |
| *Controlling for household income* | | | | |
| Indirect effect | -1.443e-5 | 9.582e-6 | 0.132 | [-3.273e-5 3.336e-6] |
| Total effect | 8.937e-5 | 7.535e-5 | 0.236 | [-5.849e-5 2.363e-4] |
| *Controlling for parental education* | | | | |
| Indirect effect | -2.180e-5 | 1.044e-5 | 0.037 | [-4.243e-5 -2.968e-6] |
| Total effect | 6.995e-5 | 7.521e-5 | 0.355 | [-7.671e-5 2.176e-5] |
| **Main model (Year 2 follow-up)** | | | | |
| Indirect effect | -8.190e-5 | 1.981e-5 | < 0.001 | [-1.243e-4, -4.440e-5] |
| Total effect | -2.856e-4 | 1.150e-4 | 0.004 | [-5.178e-4, -4.681e-5] |
| *Controlling for sex and age* | | | | |
| Indirect effect | -6.008e-5 | 1.463e-5 | < 0.001 | [-9.068e-5, -3.539e-5] |
| Total effect | -2.650e-4 | 9.143e-5 | 0.004 | [-4.442e-4, -9.474e-5] |
| *Controlling for household income* | | | | |
| Indirect effect | -3.155e-5 | 1.057e-5 | 0.003 | [-5.649e-5, -1.473e-5] |

| | | | | |
|---|---|---|---|---|
| Total effect | -1.934e-4 | 9.174e-5 | 0.035 | [-3.620e-4, -1.804e-5] |
| *Controlling for parental education* | | | | |
| Indirect effect | -3.629e-5 | 1.171e-5 | 0.002 | [-6.275e-5, -1.630e-5] |
| Total effect | -2.072e-4 | 9.133e-5 | 0.023 | [-3.841e-4, -3.251e-5] |

### D. Mediation model with reward sensitivity as the mediator

To rule out the possibility that it is reward sensitivity mediating the relationship between screen time and inhibitory control network, we also tested a regression model using screen-activity proportion as the independent variable, BAS as the mediator, and the strength of the fronto-striatal connectivity as outcome. This alternative model did not show a statistically significant mediation effect (indirect effect: $B$ = 3.477e-4, $SE$ = 3.687e-4, $p$ = 0.346, 95% $CI^{5,000}$ bootstrap = [-4.054e-4, 0.001]; total effect: $B$ = -0.008, $SE$ = 0.002, $p$ < 0.001, 95% $CI^{5,000}$ bootstrap = [-0.012, -0.003]).

### E. Association between screen exposure and the subdivisions of the striatum.

Previous studies have reported that different subdivisions of the striatum serve distinct functions on forming a behavior, such that the ventral striatum (i.e., Nucleus Accumbens) is playing a role in voluntary behavior but the dorsal striatum (i.e., Caudate and Putamen) is related to involuntary addiction (Everitt et al., 2013; Zhou et al., 2018). Therefore, we examined the association between children's baseline year Screen-Activity Proportion (SAP) score to two separate years of the neural connectivity data between the Frontoparietal network to each subdivision of the striatum (**Figure S2**). The results showed that there were stronger significant differences in the connectivity to both dorsal striatal divisions, Caudate and Putamen, in the year 2 (**Table S3**).

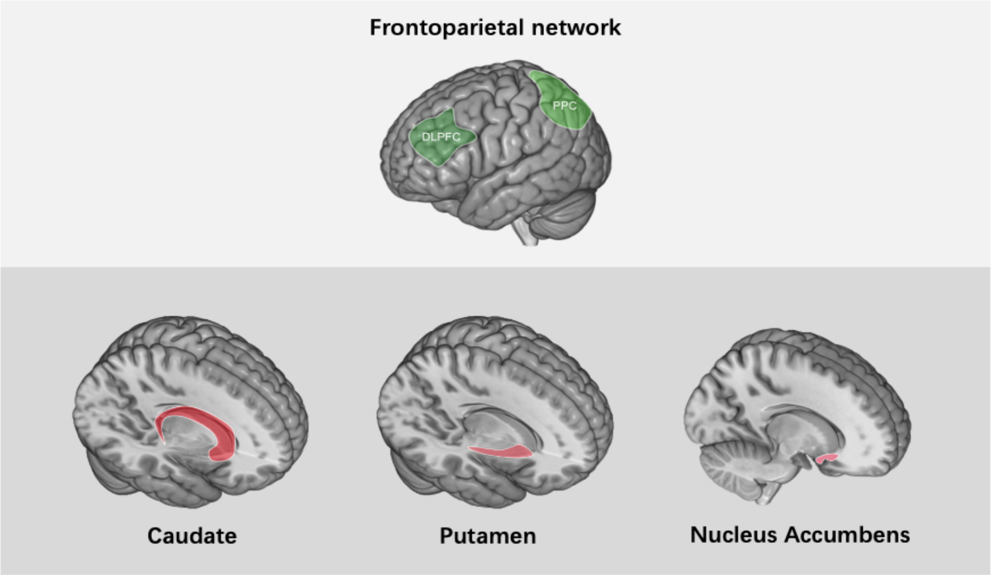


***Figure S2.*** *Illustrations of the ROIs examined in the subdivision connectivity test.*

***Table S3.*** *The results of the association between screen exposure and the subdivisions of the striatum*

| | ***r*** | **CI$^{50,000}$ bootstrap** | ***$BF_{10}$*** |
|---|---|---|---|
| **Baseline year** | | | |
| Caudate | -0.0273 | [-0.0481, -0.0066] | 0.535 |
| Putamen | -0.0215 | [-0.0423, -0.0004] | 0.173 |
| Nucleus Accumbens | -0.0246 | [-0.0456, -0.0039] | 0.303 |
| **Year 2 follow-up** | | | |
| Caudate | -0.0597 | [-0.0902, -0.0292] | 15.738** |
| Putamen | -0.0710 | [-0.1010, -0.0414] | 245.716*** |
| Nucleus Accumbens | -0.0257 | [-0.0560, 0.0047] | 0.070 |

*Caudate: FPN-Caudate connectivity; Putamen: FPN-Putamen connectivtiy; Nucleus Accumbens: FPN-Nucleus Accumbens connectivity.*

** BF10 > 3;** BF10 > 10; *** BF10 > 30.*

### F. Zero-order correlations of the main variables

***Table S4.*** *Zero-order correlation between screen exposure, reward sensitivity, and the inhibitory control network in the baseline year with Pearson's r (p-value), 95% $CI^{5,000}$ bootstrap, and $BF_{10}$.*

| ***Variables*** | ***Screen exposure*** | ***Reward sensitivity*** |
|---|---|---|
| ***Reward sensitivity*** | 0.164 (<0.001)<br>95% $CI^{5,000}$ bootstrap<br>[0.142, 0.185]<br>$BF_{10}$ = 6.815e+44 | - |
| ***FPN-Striatum*** | -0.040 (<0.001)<br>95% $CI^{5,000}$ bootstrap<br>[-0.060, -0.019]<br>$BF_{10}$ = 13.919 | 0.004 (0.720)<br>95% $CI^{5,000}$ bootstrap<br>[-0.018, 0.027]<br>$BF_{10}$ = 0.030 |

***Table S5.*** *The results of correlation between screen exposure and FPN-striatal connectivity with controlling nuisance regressors.*

| | ***r*** | ***p*** | ***95% $CI^{5,000}$ bootstrap*** |
|---|---|---|---|
| **Baseline year** | | | |
| **Screen exposure x FPN-Striatum** | -0.040 | <0.001 | [-0.060, -0.019] |
| *Controlling for sex and age* | -0.041 | <0.001 | [-0.062, -0.021] |
| *Controlling for household income* | -0.016 | 0.154 | [-0.037, 0.005] |
| *Controlling for parental education* | -0.021 | 0.060 | [-0.042, -4.444e-5] |
| **Year 2 follow-up** | | | |
| **Screen exposure x FPN-Striatum** | -0.078 | <0.001 | [-0.111, -0.046] |
| *Controlling for sex and age* | -0.079 | <0.001 | [-0.111, -0.046] |
| *Controlling for household income* | -0.057 | <0.001 | [-0.090, -0.024] |
| *Controlling for parental education* | -0.052 | 0.001 | [-0.085, -0.019] |

**G. Associations between screen exposure, brain, and behavioral indices of inhibitory function**

Associations between screen exposure, brain, and behavioral indices of inhibitory function were provided for validating the effect of screen exposure and resting-state FPN-Striatum functional connectivity anchor on inhibitory control ability. The behavioral indices of inhibitory function included the performance in the Flanker Inhibitory Control and Attention Test (*abcd_tbss01*, age-corrected), the Monetary Incentive Delay Task (*abcd_mid02*, total earn), the Stop Signal Task (*abcd_sst02*, stop-signal reaction time), ADHD trait (abcd_cbcls01, t-score) and the Cash Choice Task (*cct01*).

**Flanker Inhibitory Control and Attention Test.** Flanker Inhibitory Control and Attention Test. The flanker task (Eriksen & Eriksen, 1974) was designed for measuring inhibitory control and visual attention function. The NIH Toolbox Flanker Inhibitory Control and Attention Test, Ages 8-11 v2.0, was used in the ABCD study. Participants have to finish two blocks of the Flanker task, a block with child-friendly fish stimuli and a block with a standard arrow. Participants were asked to point out the left or right orientation of a central stimulus while inhibiting a distraction from the flankers on each side. In some trials, the orientation of the central stimulus was congruent with the orientation of the flankers, while in others it was incongruent. During incongruent trials, more inhibitory control effort is recruited. The current analysis utilized the age-corrected standard score with a higher score indicating a better control ability (Zelazo et al., 2013).

**Monetary Incentive Delay Task (MID).** The MID task (Everitt et al. 2008) is a widely used approach to measure neural mechanisms of reward processing. The current analysis only included the behavioral performance (i.e., total earn) as an inhibitory control index, which earns more indicating performs better. The ABCD version of the MID task begins with an incentive cue with five possibilities (Win $5.0, Win $0.2, Lose $0.2, Lose $5.0, or $0 - no money at stake), a delay followed by a response phase is designed to measure the individual ability of anticipation and inhibitory control. Of the total 100 trials, 40 trials were assigned as reward anticipation, 40 as loss anticipation, and 20 as no money anticipation trials.

**Stop Signal Task (SST).** The SST (Logan, 1994) measures impulsivity and impulse control ability. It requires participants to withhold their motor responses to a “Go” stimulus when an unpredictable “Stop” is presented. Participants were instructed to respond as quickly and accurately as possible when a “Go” signal is presented. They were also instructed to withhold the motor response when a delayed “Stop” signal is presented. If participants are able to withhold the motor response while “Stop” is shown, the trial is counted as successful inhibition, whereas if participants still respond, the trial is counted as unsuccessful inhibition. Of the total 180 trials in each run of the task, 30 of the trials (16.67%) were “Stop” trials. To ensure that there were about 50% successful and 50% unsuccessful inhibition trials, a tracking algorithm varied the interval between the “Go” signal onset and the “Stop” signal onset. The stop-signal reaction time (SSRT) was calculated by subtracting the average of the stop-signal delay from the average of the go response time. A shorter SSRT corresponds to better inhibitory control (Logan & Cowan, 1984; Logan et al., 2014). More details about the design of MID and SST are available in Casey et al. (2018).

**ADHD trait.** The ADHD Scale (t-score) scale of the child behavior checklist (CBCL) was used as the measure of ADHD traits, with a higher score indicating more behavioral problems (Achenbach & Edelbrock, 1991; Owens et al., 2021).

**Cash Choice Task.** In the task, participants were instructed to pretend a kind person wanted to give them some money. Participants have to make a binary decision: Would you rather have $75 in three days or $115 in 3 months? (1 = $75 in three days; 2 = $115 in 3 months).

***Table S6.*** *The results of the association between screen exposure, brain, and behavioral indices of inhibitory function with r and [$CI^{50,000}$bootstrap].*

| | ***SAP (baseline)*** | ***FPN-Striatum (baseline)*** | ***FPN-Striatum (2 year follow-up)*** |
|---|---|---|---|
| **Sample size** | 8,324 | 8,324 | 3,891 |
| **Baseline year** | | | |
| Flanker Task | -0.0904*** [-0.1114, -0.0690] | 0.0151 [-0.0084, 0.0384] | - |
| MID | -0.0581*** [-0.0801, -0.0366] | 0.0394*** [0.0178, 0.0609] | - |
| SSRT | 0.0022 [-0.0194, 0.0238] | -0.0247* [-0.0478, -0.0018] | - |
| ADHD | 0.0778*** [0.0554, 0.0997] | -0.0207 [-0.0527, 0.0114] | - |
| Cash Choice | -0.0285** [-0.0503, -0.0070] | 0.0047 [-0.0173, 0.0270] | - |
| ***Year 2 follow-up*** | | | |
| Flanker Task | [a]-0.1152*** [-0.1417, -0.0887 | [a]0.0164 [-0.0110, 0.0441] | 0.0447* [0.0119, 0.0777] |
| MID | [b]-0.0314* [-0.0581, -0.0050] | [b]0.0116 [-0.0158, 0.0389] | 0.0035 [-0.0276, 0.0351] |
| SSRT | 0.0250 [-0.0066, 0.0566] | -0.0060 [-0.0377, 0.0256] | 0.0281 [-0.0073, 0.0627] |
| ADHD | [c]0.0901*** [0.0624, 0.1173] | [c]-0.007 [-0.0363, 0.0223] | -0.0306 [-0.0635, 0.0023] |

*ADHD: Attention Deficit Hyperactivity Disorder; MID: Total Earn in the Monetary Incentive Delay task; SAP: Screen-Activity Proportion score; SSRT: Stop Signal Reaction Time*
*[a]N = 5570; [b]N = 5486; [c]N = 5686. *p < .05; **p < .01; ***p < .001.*

### Supplementary References


Achenbach, T. M., & Edelbrock, C. (1991). Child behavior checklist. Burlington (Vt), 7, 371-392.

Casey, B., Cannonier, T., Conley, M. I., Cohen, A. O., Barch, D. M., Heitzeg, M. M., . . . Garavan, H. (2018). The adolescent brain cognitive development (ABCD) study: imaging acquisition across 21 sites. *Developmental cognitive neuroscience, 32,* 43-54.

Eriksen, B. A., & Eriksen, C. W. (1974). Effects of noise letters upon the identification of a target letter in a nonsearch task. *Perception & psychophysics, 16*(1), 143-149.

Everitt, B. J., Belin, D., Economidou, D., Pelloux, Y., Dalley, J. W., & Robbins, T. W. (2008). Neural mechanisms underlying the vulnerability to develop compulsive drug-seeking habits and addiction. *Philosophical Transactions of the Royal Society B: Biological Sciences, 363*(1507), 3125-3135.

Everitt, B. J., & Robbins, T. W. (2013). From the ventral to the dorsal striatum: devolving views of their roles in drug addiction. *Neuroscience & Biobehavioral Reviews, 37*(9), 1946-1954.

Logan, G. D. (1994). On the ability to inhibit thought and action: A users' guide to the stop signal paradigm.

Logan, G. D., & Cowan, W. B. (1984). On the ability to inhibit thought and action: A theory of an act of control. *Psychological review, 91*(3), 295.

Logan, G. D., Van Zandt, T., Verbruggen, F., & Wagenmakers, E.-J. (2014). On the ability to inhibit thought and action: general and special theories of an act of control. *Psychological review, 121*(1), 66.

Owens, M. M., Allgaier, N., Hahn, S., Yuan, D., Albaugh, M., Adise, S., . . . Potter, A. (2021). Multimethod investigation of the neurobiological basis of ADHD symptomatology in children aged 9-10: baseline data from the ABCD study. *Translational psychiatry, 11*(1), 1-11.

Zelazo, P. D., Anderson, J. E., Richler, J., Wallner-Allen, K., Beaumont, J. L., & Weintraub, S. (2013). II. NIH Toolbox Cognition Battery (CB): Measuring executive function and attention. *Monographs of the Society for Research in Child Development, 78*(4), 16-33.

Zhou, F., Zimmermann, K., Xin, F., Scheele, D., Dau, W., Banger, M., ... & Becker, B. (2018). Shifted balance of dorsal versus ventral striatal communication with frontal reward and regulatory regions in cannabis-dependent males. *Human brain mapping, 39(12),* 5062-5073.
